# Systemic aneuploidization events drive phenotype switching in *Saccharomyces cerevisiae*

**DOI:** 10.1101/2021.06.16.448724

**Authors:** Lydia R. Heasley, Juan Lucas Argueso

## Abstract

How cells leverage their phenotypic potential to adapt and survive in a changing environment is a complex biological problem, with important implications for pathogenesis and species evolution. One particularly fascinating adaptive approach is the bet hedging strategy known as phenotype switching, which introduces phenotypic variation into a population through stochastic processes. Phenotype switching has long been observed in species across the tree of life, yet the mechanistic causes of switching in these organisms have remained difficult to define. Here we describe the causative basis of colony morphology phenotype switching which occurs among cells of the pathogenic isolate of *Saccharomyces cerevisiae*, YJM311. From clonal populations of YJM311 cells grown in identical conditions, we identified colonies which displayed altered colony architectures, yet could revert to the wild-type morphology after passaging. Whole genome sequence analysis revealed that these variant clones had all acquired whole chromosome copy number alterations (*i.e.,* aneuploidies). Cumulatively, the variant clones we characterized harbored an exceptional spectrum of karyotypic alterations, with individual variants carrying between 1 and 16 aneuploidies. Most variants harbored unique collections of aneuploidies, indicating that numerous distinct karyotypes could manifest in the same morphological variation. Intriguingly, the genomic stability of these newly aneuploid variant clones modulated how often cells reverted back to the wild-type phenotypic state. We found that such revertant switches were also driven by chromosome missegregation events, and in some cases occurred through a return to euploidy. Together, our results demonstrate that colony morphology switching in this yeast strain is driven by stochastic and systemic aneuploidization events. These findings add an important new perspective to our current understanding of phenotype switching and bet hedging strategies, as well as how environmental pressures perpetuate organismal adaption and genome evolution.

## Introduction

The ability to adapt to fluctuating environmental conditions is critical to the survival of organisms. Many adaption strategies rely on the regulated activities of environmental response pathways, which enable a cell to sense an environmental change, and induce a programmed response such as to alter its phenotype accordingly^1^. Alternatively, cells may stochastically shift between phenotypic states, even in the absence of environmental stimuli. This process, called phenotype switching, generates phenotypic heterogeneity within a clonal population, and can produce subpopulations of cells better equipped to withstand an acute environmental fluctuation^2,3^. For this reason, stochastic phenotype switching has been proposed to function as a bet hedging strategy, as switching has been found to increase the overall adaptive potential of a population^3–6^.

Stochastic switches in microbial colony morphology (CM) are iconic examples of phenotype switching. Among the best described are the switches displayed by the opportunistic pathogen *Candida albicans*^7,8^. *C. albicans* cells normally form smooth colonies, yet can stochastically switch to form a repertoire of highly differentiated colony structures^8^. Such alterations in colony structure are thought to promote population survival in fluctuating environments by enabling cell specialization, resource sharing, and cooperative growth^9,10^. Similar morphological switches have also been observed in the pathogen *Cryptococcus neoformans* as well as in pathogenic isolates of the budding yeast *Saccharomyces cerevisiae*^11,12^.

CM switching has been described for numerous species, yet the biological mechanisms which underlie stochastic switches are less well understood^4^. Insights may perhaps be gleaned from recent works defining the developmentally programmed CM responses that are induced by environmental stimuli^9,10,13,14^. One such developmental program, referred to as the colony morphology response (CMR), is activated by changes in the availability of key nutrient factors such as carbon and nitrogen in the environment^10^. While the triggering environmental cue and the resulting morphology of CMR-activated colonies varies among isolates, the molecular underpinnings of this response are highly conserved^10^. The cellular regulation of CMR activity has been extensively described in YJM311, a pathogenic isolate of *S. cerevisiae*^15,16^. YJM311 cells activate the CMR in response to glucose deprivation. On glucose-rich media (2.0% glucose, YP^2.0%^) CMR activity is low, and cells form simple, smooth colonies (Fig. 1A, left). On glucose-poor media (0.5% glucose, YP^0.5%^) the CMR is induced, and cells form colonies with a complex and ruffled architecture (Fig. 1A, right). The diploid genome of YJM311 is highly heterozygous (>40,000 heterozygous single nucleotide polymorphisms, hetSNPs) and quantitative trait locus analysis has identified many genomic loci that contribute to expression of the CMR^15,17^. Collectively, these loci function in the glucose responsive cAMP/PKA signaling pathway and the downstream stress response and flocculation pathways^14,15^. Additionally, these studies have also revealed that allelic variation present at CMR-regulatory loci in the YJM311 genome dramatically modulates CMR activity in cells^15^.

**Figure 1.**
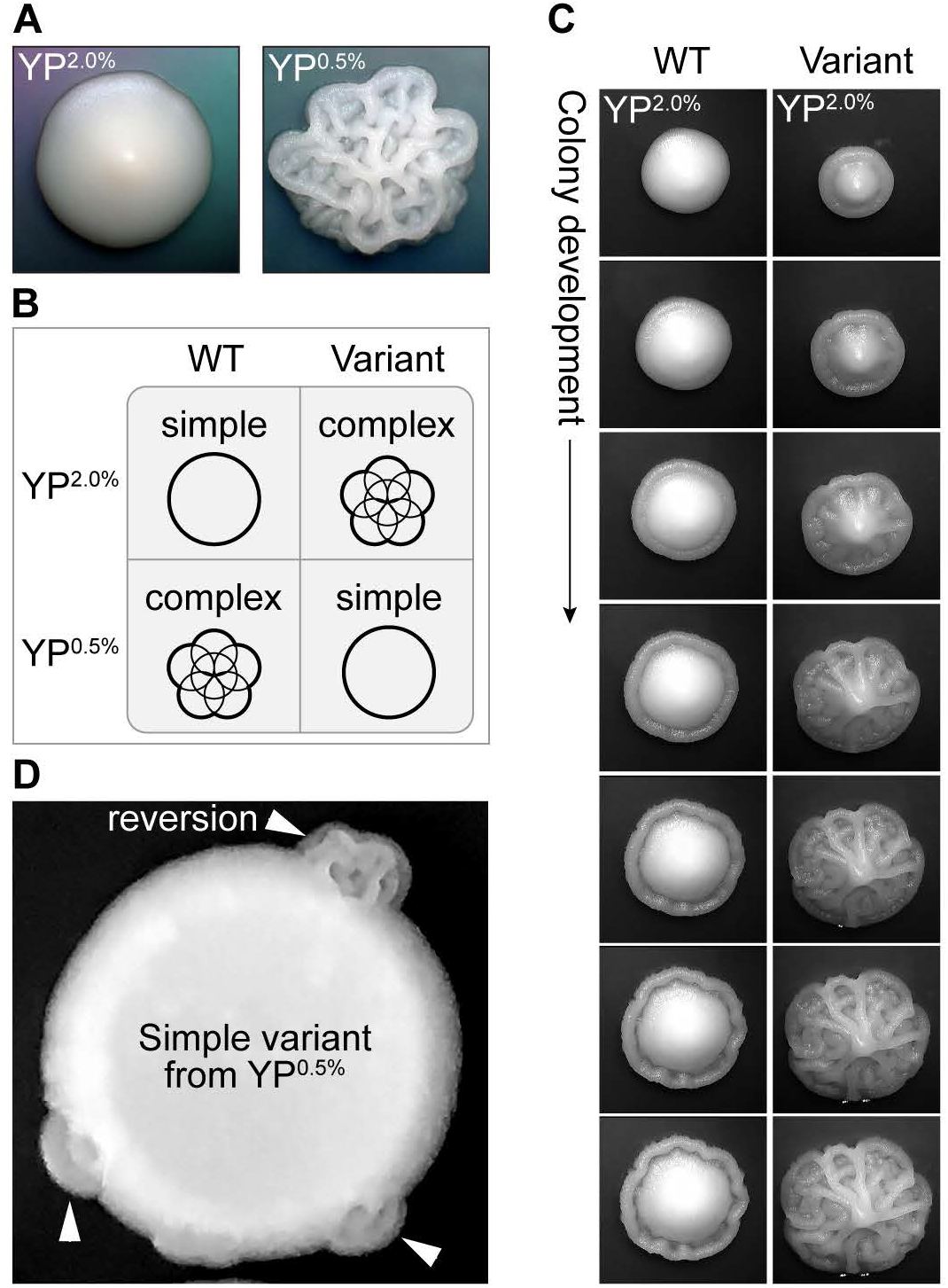
Phenotypic variants spontaneously arise in populations of YJM311. **A.** Representative images of YJM311 colony morphologies grown on YP^2.0%^ (left) YP^0.5%^ (right). **B.** A cartoon schematic outlining the morphological designations of wild-type (WT) and variant colonies on either YP^2.0%^ or YP^0.5%^. **C.** Representative time-course images of a 2-day old WT and complex variant colony developing over 24 hours on YP^2.0%^. **D.** An image depicting a cell spot of a simple variant isolated from YP^0.5%^; the smooth variant phenotype persists throughout the majority of the spot, but regions denoted by white arrowheads have reverted back to the WT morphology (*i.e.,* complex).

The morphological attributes of CMR-activated colonies are likely to be clinically relevant, as they promote invasive growth, substrate adherence, and biofilm formation^18^. Indeed, switching to a complex morphology has been shown to promote virulence and drug resistance in *C. albicans*^8,19^. This association with virulence, combined with the numerous examples of CM switching among pathogenic microbes^7,20^, underscores the importance of understanding how these switches operate, and how they contribute to the adaptive potential of microbial populations. In this study, we describe our investigation of stochastic CM phenotype switching displayed by YJM311 cells, and our discovery that genomic copy number alterations resulting from systemic aneuploidization events are dominant modulators of phenotypic plasticity.

## Results

### Phenotypic variants spontaneously arise in clonal populations

When clonal populations of YJM311 cells are plated onto glucose-rich media (YP^2.0%^), the majority of cells form simple, smooth colonies (Fig. 1A, left). However, we observed that on YP^2.0%^, ~1 in 1000 colonies displayed a distinctly complex morphology (average variant frequency, 1.29×10^−3^) (Fig. 1B, Table S1). This complex morphology was evident early in colony development and became progressively more distinguishable as colonies expanded (Fig. 1C). Such phenotypic heterogeneity was also apparent among YJM311 cells grown on YP^0.5%^ plates, a condition in which the majority of cells form complex colonies as a result of CMR activation. When examining colonies grown on YP^0.5%^, we identified simple variants, also at a frequency of ~1 in 1000 colonies (average variant frequency, 1.05×10^−3^) (Fig. 1B, Table S1). We verified that these phenotypic alterations did not result from local changes in glucose concentration by spotting variant colonies to new plates. In many cases, the spotted population displayed the variant phenotype, but some spots yielded sectors which had reverted to the wild-type (WT) phenotype (Fig. 1D, arrowheads), demonstrating that this variant phenotypic state was transient and reversible. Together, these observations suggested that stochastic heterogeneity in CM arose in populations of YJM311 cells, possibly through a switch-like mechanism.

### Phenotypic variants harbor elaborate configurations of de novo copy number alterations

We next sought to explore the causative basis of this phenotypic variation. Similar to CM switches described in other fungal systems^7,8^, the frequency at which we recovered variants was higher than has been reported for the occurrence of somatic mutations in *S. cerevisiae*. Yet, because these variant phenotypes persisted through passaging, we investigated the possibility that they could have arisen from *de novo* mutations. We performed a preliminary characterization of variant genomes using pulsed-field gel electrophoresis (PFGE), a molecular karyotyping approach that resolves yeast chromosomes by size to enable detection of gross chromosomal rearrangements and copy number alterations. We karyotyped 5 matched pairs of WT and simple variant colonies, each pair of which had been isolated from independent populations of YJM311 cells grown on YP^0.5%^. Whereas WT colonies displayed no obvious karyotypic changes relative to each other or the parental YJM311 strain, all five simple variants displayed alterations in chromosomal band intensity or pattern that were indicative of *de novo* structural variations (SVs) (Fig. 2A, arrowheads). To expand on this preliminary result, we collected a set of WT clones (20 clones), as well as complex (22 clones isolated from YP^2.0%^) and simple (26 clones isolated from YP^0.5%^) variant clones for comprehensive genomic analysis using whole genome sequencing (WGS).

**Figure 2.**
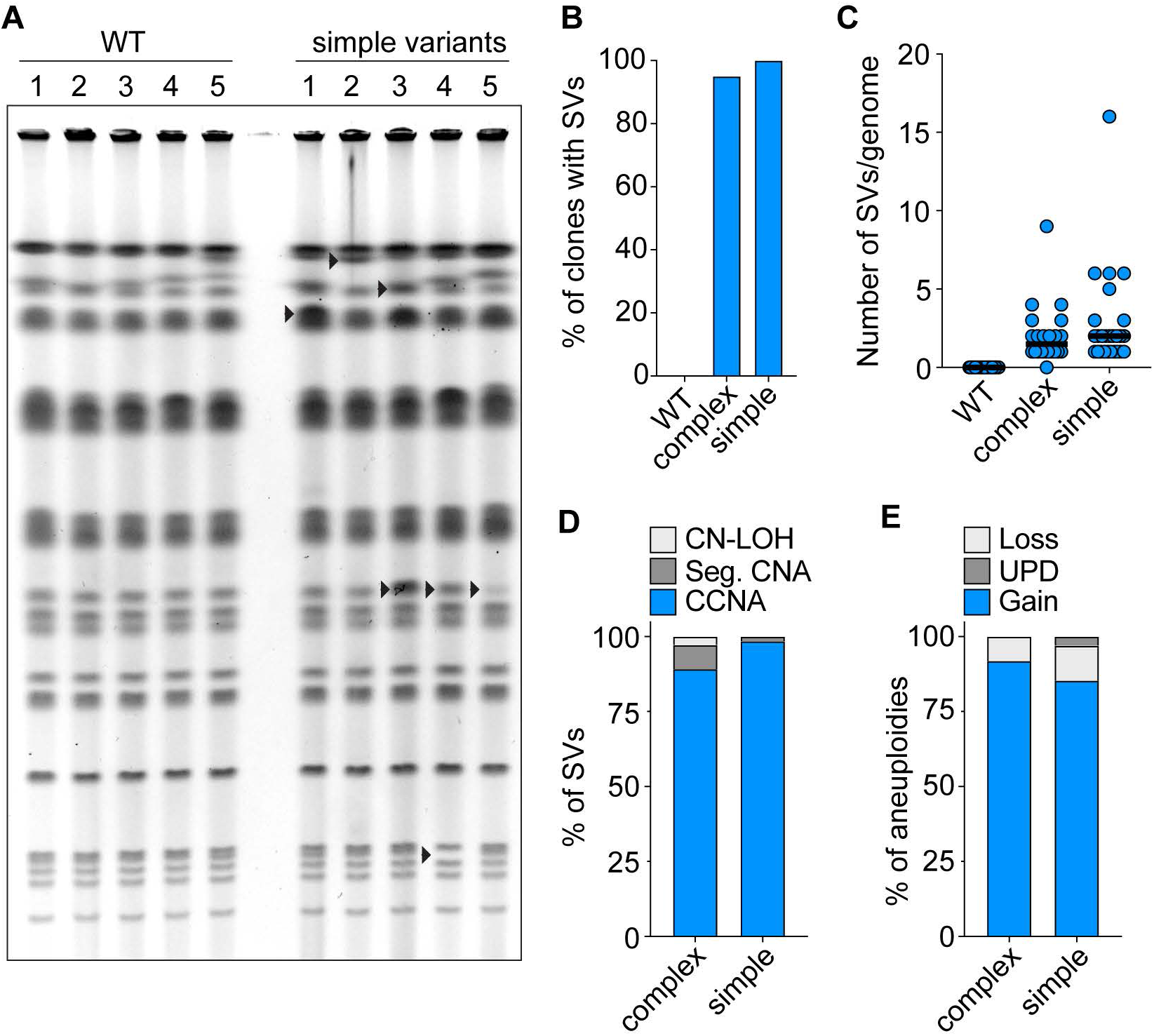
Variant clones harbor numerous *de novo* SVs. **A.** The molecular karyotypes of WT and simple variants recovered from five independent cultures (1-5). Arrowheads denote *de novo* karyotypic alterations. **B.** The percentage of WT or variant clones harboring *de novo* SVs. **C.** The number of SVs present in WT and variant genomes. Each blue circle denotes an individual clone. Black bar denotes the median number of SVs harbored by WT, complex variant, or simple variant clones (WT, 0; complex and simple, 2). **D.** The percentage of SVs that constitute copy-neutral loss-of-heterozygosity (CN-LOH), segmental copy number alteration (Seg. CNA), or whole chromosome copy number alterations (CCNAs). **E.** The percentage of CCNAs detected in variant genomes that constituted a chromosomal loss, uniparental disomy (UPD), or chromosomal gain. For all data presented, aneuploidy of Chr1^cir^ has been excluded from analysis.

We expected that all WT clones would harbor the parental YJM311 karyotype, yet surprisingly, our WGS analysis revealed that 20.0% (4/20) were monosomic for chromosome 1 (Chr1). In a separate and forthcoming study in which our group characterized the fully phased diploid genome of YJM311, we discovered that one homolog of Chr1, which we term Chr1^cir^, is a circularized chromosome (Fig. S1). Due to its circular nature, this chromosome cannot be resolved by PFGE, and was therefore precluded from our preliminary karyotypic analysis (Fig. 2A). Analysis of sequencing data generated from our YJM311 stock established that it is a mosaic population consisting of euploid cells and cells lacking Chr1^cir^ (Table S2). Indeed, Chr1^cir^ was lost in all four monosomic WT clones, suggesting that these clones were derived from the pre-existing monosomic cells in the stock population. We also found that some euploid WT clones displayed intra-clonal mosaicism for Chr1^cir^ loss, indicating that although the cell which founded the colony had been euploid, Chr1^cir^ had been lost in a subpopulation of cells as the colony expanded (Table S2). Together, these findings suggest that Chr1^cir^ may be an intrinsically unstable chromosome, likely due to its circular structure^21,22^. Among the collection of complex and simple variant clones, 4.5% (1/22) and 15.3% (4/26) had also lost Chr1^cir^, respectively. However, because cells lacking Chr1^cir^ display the WT CM, monosomy of Chr1 is unlikely to be linked to the CM phenotype switching we were investigating. For this reason, we excluded loss of Chr1^cir^ from our subsequent genomic analyses of WT and variant clones. Besides aneuploidy of Chr1^cir^, we detected no other karyotypic alterations in the genomes of WT clones (Fig. 2B). In striking contrast, however, we detected numerous *de novo* SVs in the genomes of variant clones. Overall, 95% of complex clones and 100% of simple clones had acquired new SVs (Fig. 2B), with individual clones harboring between 1-16 new SVs per genome (median, 2 SVs/ genome) (Fig. 2C, Table S3). Interestingly, the vast majority of these SVs were whole chromosome copy number alterations (CCNAs) (*i.e.,* aneuploidies), and most often constituted chromosomal gains (Fig. 2D-E).

While most variants harbored unique karyotypic configurations (Fig. 3A-B), we also identified specific aneuploidies that were enriched in either the complex or simple sets of variant clones (Fig. 3C). For example, gains of Chr13 were more often carried by complex clones than simple clones (13/22 complex variants; 3/26 simple variants) (Fig. 3A,C). Similarly, gains of Chr15 were more often harbored by simple clones than by complex clones (13/26 simple variants, 1/22 complex variants) (Fig. 3B, C). Several variant clones had acquired only an additional copy of Chr13 (6/22 complex variants) or Chr15 (6/26 simple variants), indicating that while not necessary, trisomy of Chr13 or Chr15 is sufficient to induce a CM switch in YJM311. A relatively smaller percentage of clones harbored segmental copy number alterations (Seg. CNAs) (10.4%, 5/48 variant clones) (Fig. 2D). Of these, four harbored non-reciprocal translocations which altered the copy number of specific chromosomal regions (Fig. S2, Table S3). Collectively, these results indicated that *de novo* copy number alterations, and specifically CCNAs, were likely the causative source of CM phenotype switching in YJM311.

**Figure 3.**
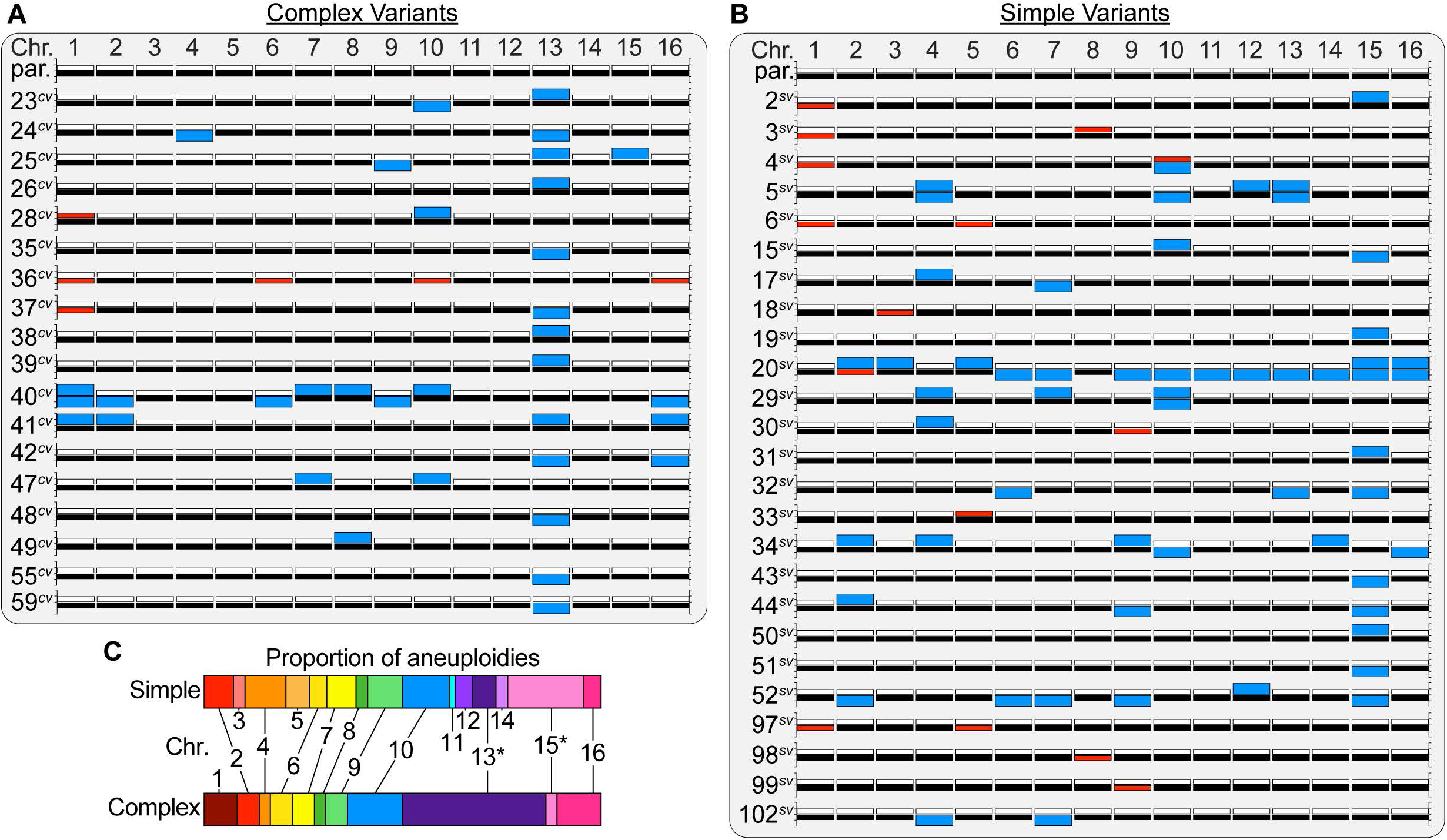
The genomes of variant clones are dramatically altered by aneuploidization. **A-B.** Karyotypes of the YJM311 parent and all complex and simple variants harboring *de novo* CCNAs. For each chromosome, white bars represent homolog ‘A’ and black bars represent homolog ‘B’. Blue bars denote a chromosomal gain, red bars denote a chromosomal loss. **C.** The proportion of total CCNAs that affected each chromosome in simple and complex variants. Asterisk denotes an aneuploidy that is significantly enriched in one class of variant clones relative to the other.

Given that the sequenced variant clones were isolated after only ~30 generations of growth (see Methods), the degree to which their genomes had been altered by CCNAs suggested that clones harboring multiple aneuploidies had likely acquired them simultaneously from a systemic error in chromosome segregation^23,24^. Indeed, most CCNAs were detected at discrete copy numbers, indicating that they did not arise continuously during colony expansion. However, we also considered the possibility that YJM311 might exhibit chromosomal instability (CIN), a mutator phenotype that results in elevated rates of chromosome loss^25^. To determine whether YJM311 cells displayed CIN, we used a counter-selectable plating assay and fluctuation analysis to calculate the rate at which cells lost one homolog of Chr5^23,26^. In the diploid YJM311 background, we deleted the counter-selectable marker *CAN1* from its endogenous locus on the left arm of one homolog (Chr5^A^) and introduced a second copy of *CAN1* onto the right arm of the other homolog (Chr5^B^)(Fig. S3)^23^. This parental strain was sensitive to the toxic arginine analog canavanine, but cells that lost Chr5^B^ were resistant and could form colonies on media containing canavanine^26^. We calculated the average rate at which cells lost Chr5^B^ to be 1.04×10^−5^ (Table S4), a value consistent with published rates of chromosome loss in *S. cerevisiae*^23,27–29^. Since YJM311 did not show elevated rates of chromosome loss relative to other laboratory strains, we conclude that a CIN phenotype is unlikely to drive the pervasive aneuploidy we observed in the genomes of variant YJM311 colonies.

### Copy number alterations, more often than allelic identity, underlie phenotypic variation

In YJM311, allelic differences at multiple genomic loci have been found to greatly modulate expression of the CMR^15^. As such, we had anticipated that some variant clones would harbor *de novo* tracts of copy neutral loss-of-heterozygosity (LOH) which would alter the allelic identity of CMR-linked genes. Yet, of the 48 variant genomes we examined, only one clone harbored a copy neutral LOH event which impacted the genotype of the CMR-linked loci *YAK1* and *CYR1* (4^*SV*^, Fig. 3B). This simple variant had acquired uniparental disomy (UPD) of Chr10, a type of CCNA that results in copy-neutral whole chromosome LOH through loss of one homolog and gain of the other^30,31^. Recovery of this variant demonstrated that copy neutral LOH is, at least in this case, sufficient to induce phenotypic switching. However, the results from our genomic analysis suggested that copy number alterations represented a prevailing mode of phenotypic variation.

We considered the possibility that variant clones might harbor CCNAs of chromosomes carrying particular alleles of CMR regulatory genes. We first tested this premise by investigating complex variant clones that had gained only an extra copy of Chr13, which carries the gene *MSN2*, a stress-responsive transcriptional activator and major CMR regulatory locus^15,32,33^. We determined whether these clones displayed a bias for having acquired a specific homolog of Chr13 (Chr13^A^ or Chr13^B^) and found that both Chr13^A^ and Chr13^B^ had been gained at equal frequencies (Table S3). Additionally, when we examined the colony morphologies of variants carrying an extra copy of either Chr13^A^ or Chr13^B^, we found that while there were modest differences in colony architecture, both variants produced obviously complex colonies on YP^2.0%^ (Fig. 4A). We conducted a similar analysis of simple variant clones harboring trisomies of Chr15, which carries the CMR-linked gene *SFL1,* a transcriptional repressor of flocculation and invasive growth^14,15,34^. Again, we found that these clones carried extra copies of either Chr15^A^ or Chr15^B^, with no apparent bias for one homolog over the other (Table S3). Clones harboring a gain of either Chr15^A^ or Chr15^B^ both formed smooth colonies on YP^0.5%^, though these morphologies were slightly distinct from one another (Fig. 4B). As a final test, we considered a case in which the ploidy of a chromosome had been reduced to monosomy, which would result in LOH at all loci present on the chromosome. We analyzed clones that had become monosomic for Chr5, which carries *FLO8*, another key effector of the CMR^15^. There were only three clones in our variant collection harboring only monosomy of Chr5, yet we found that two had lost Chr5^A^, while the third had lost Chr5^B^ (Table S3). Moreover, clones monosomic for either homolog of Chr5 formed simple colonies on YP^0.5%^ that were indistinguishable from one another (Fig. 4C). Taken together, these results indicate that although allelic variation at CMR-linked loci can modulate CM, the molecular basis of CM phenotype switching in YJM311 occurs primarily through gene dosage effects resulting from aneuploidization.

**Figure 4.**
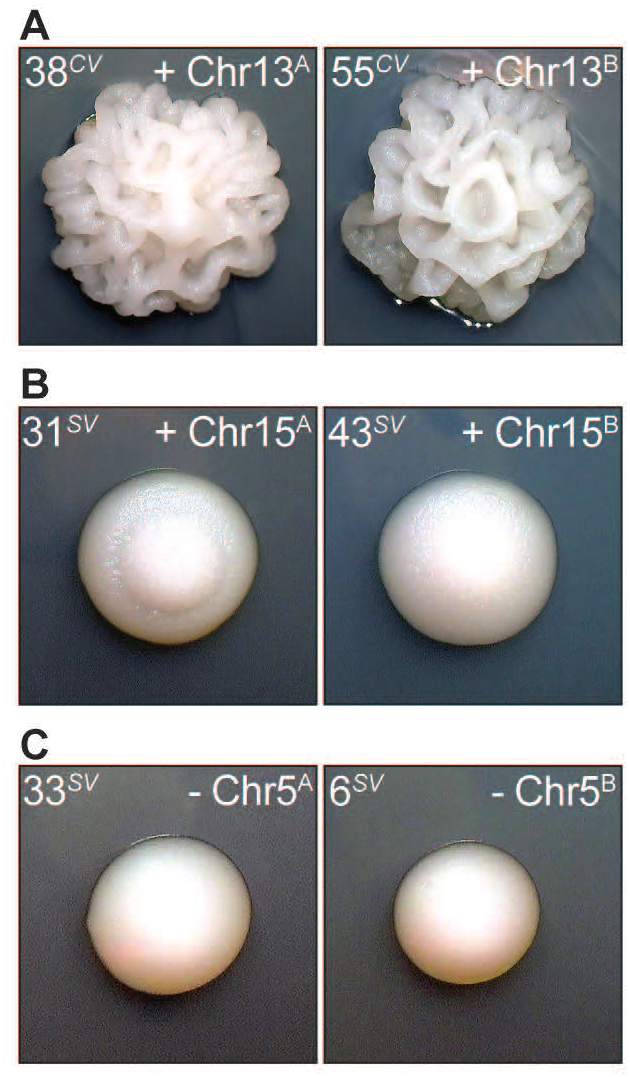
Copy number variations, not allelic identity, are dominant drivers of phenotypic variation **A-C.** Representative images of colony morphology phenotypes displayed by the denoted variants. Colonies depicted in A. were grown on YP^2.0%^ plates. Colonies depicted in B-C were grown on YP^0.5%^ plates.

### Unique aneuploid configurations differentially modulate switching frequencies by altering genome stability

The ability to toggle between phenotypic states is a defining feature of phenotype switching. Since somatic mutations usually impart permanent and heritable genotypic changes to a cell, they have been generally dismissed as putative mechanisms of switching. Yet, because aneuploid cells can return to euploidy through subsequent chromosome missegregation events, aneuploidization represents a potentially reversible genomic alteration^5,31,35,36^. Thus, we investigated whether phenotypic switches driven by CCNAs assumed a toggle-like quality. To do so, we used a colony sectoring assay to measure the frequencies at which variant clones switched back to the WT phenotype. We performed sectoring analysis for variants harboring monosomy of Chr5 (6^*SV*^), Chr9 (99^*SV*^), or Chr3 (18^*SV*^), or trisomy of Chr10 (28^*CV*^), Chr13 (35^*CV*^), Chr15 (31^*SV*^), or both Chr4 and Chr7 (17^SV^). Cells from each variant clone were plated to the media from which they were originally isolated (YP^2.0%^ or YP^0.5%^), and colonies were screened for sectors that displayed the revertant phenotype (Fig. 5A). Interestingly, this set of aneuploid variants displayed a broad spectrum of reversion frequency. On average, clones harboring monosomic karyotypes produced revertant sectors 10-fold more often than clones harboring trisomic karyotypes. Yet, within these subsets, reversion frequencies varied between specific clones, suggesting that the genomic stability of each unique karyotypic configuration differentially modulated the frequency of phenotype switching (Fig. 5B, Table S5).

**Figure 5.**
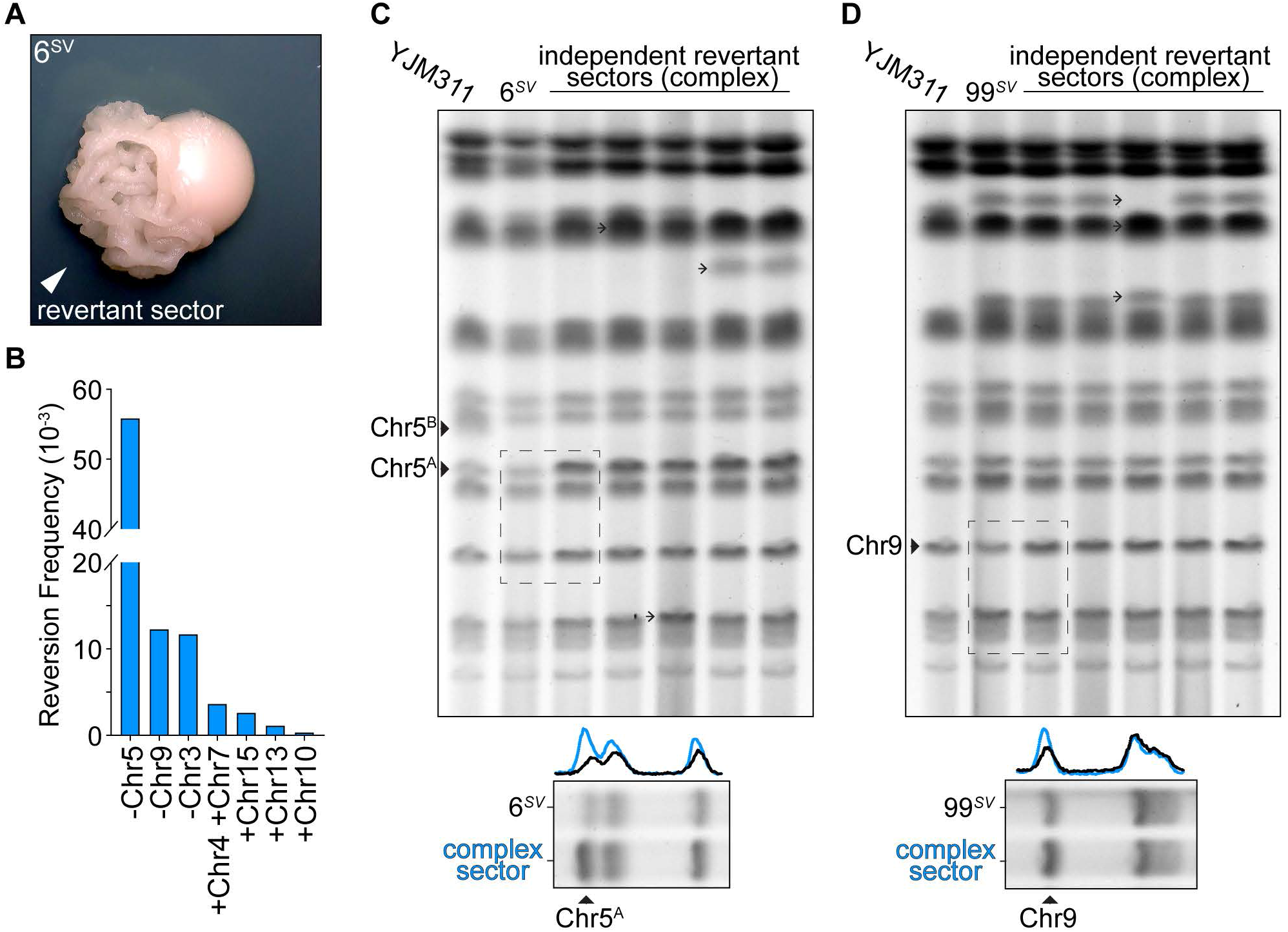
Karyotypic configurations modulate the frequency of phenotype switching. **A.** A representative image of a sectored colony from clone 6^*SV*^ (monosomy of Chr5) grown on YP^0.5%^. Arrowhead indicates the sector that has reverted back to the wild-type morphology. **B.** The frequency of revertant sectors calculated for variant isolates harboring the denoted karyotypic alterations. **C.** Top, molecular karyotypes of the parental YJM311 genome, the simple variant 6^*SV*^, and five revertant sectors recovered from independent colonies. Solid arrowheads denote the bands correlating to the two homologs of Chr5 (Chr5^A^, Chr5^B^). 6^*SV*^ harbors a loss of Chr5^B^. Barbed arrowheads denote additional SVs that arose concurrently in the genomes of revertant sectors. Bottom, band intensity tracing analysis and representative crop from the above image (dashed rectangle) confirming UPD of Chr5^A^ in revertant complex sectors. Black trace represents band intensities derived from 6^*SV*^, blue trace represents band intensities derived from a complex sector. The increased amplitude of the blue peak corresponding to Chr5^A^ correlates with an increase in copy number from 1 to 2 copies. **D.** Same as in C, but depicting karyotypic analysis of the simple variant 99^*SV*^ (monosomy of Chr9) and five revertant sectors.

We investigated the basis of these reversion events using the monosomic clones 6^*SV*^ and 99^*SV*^. We hypothesized that for these clones, reversions likely arose by a return from monosomy to UPD via a second aneuploidization event. To determine whether UPD correlated with phenotypic reversion, we performed PFGE molecular karyotype analysis for the genomes of the parental variant clone and five derivative revertant sectors (Fig. 5C-D). Indeed, PFGE band intensity analysis showed that all revertant sectors recovered from 6^*SV*^ and 99^*SV*^ had acquired UPD of Chr5 or Chr9, respectively (Fig. 5C-D, bottom). In addition to harboring UPD of Chr5 or Chr9, several revertant sectors also acquired additional new SVs (Fig. 5C-D, top, arrows), indicating that in these cases, the reversion was associated with a systemic aneuploidization event affecting other chromosomes. Together, these results demonstrated that CCNA-driven events facilitate switches between phenotypic states. Moreover, they demonstrated that the stability of these aneuploid variant genomes modulates the frequency of subsequent switching events.

## Discussion

Here we have presented data demonstrating that the CM switching behavior of a pathogenic isolate of *S. cerevisiae* is driven by aneuploidization of the genome. While often portrayed as an exclusively detrimental genomic alteration^25,37,38^, aneuploidization is also recognized as a ubiquitous adaptive strategy used by cells to survive fluctuating environmental conditions^36,39,40^. Aneuploidies are acquired at rates higher than other genomic mutations and can confer short- and long-term advantages to a cell by acutely and reversibly impacting gene expression and by enabling the acquisition of permanent adaptive mutations^35,41,42^. In fact, like phenotype switching, aneuploidization has also been suggested to function as a bet hedging strategy, as it can introduce significant phenotypic diversity into a population simply through stochastic errors in chromosome segregation^41^. Our observation that these bet hedging strategies mutually generate CM variation in *S. cerevisiae* contributes an important perspective to our understanding of how environmental adaptation can shape the genomes of organisms and the diversity of populations.

The potential impacts of aneuploidy-driven phenotype switching on population diversity are further heightened when framed within the emergent concept of systemic genomic instability (SGI)^23,24,43,44^. Recent work from our group demonstrated that aneuploidization events often impart systemic genomic changes, with the potential to impact the transmission of many, if not all, chromosomes in the cell concurrently. Consequently, these events produce a spectrum of remarkably altered karyotypes^23^, such as those harbored by the phenotypic variants characterized in the present study. Here, our results highlight how the systemic nature of aneuploidization can generate genotypically unique cells that display a similarly altered phenotype (*i.e.,* the formation of complex colonies on YP^2.0%^). This, together with the fact that the CMR is regulated by numerous genes located on multiple chromosomes^15^, explains the frequency at which we detected CM variants in populations of YJM311 cells.

Perhaps the most interesting implication of systemic aneuploidization as it relates to phenotype switching derives from our finding that the stability of the new karyotypic configurations harbored by phenotypic variants modulates the properties of future switching events. Even among variants harboring just one aneuploid chromosome, we observed notable differences in the frequencies at which cells reverted back to the WT phenotype. Thus, systemic aneuploidization not only underlies this phenotypic switch, but it also introduces plasticity into the switching behaviors displayed by genotypically unique variants. Collectively, this switching system stochastically introduces four types of diversity into a population of cells: 1) diversity in CM and the associated environmental sensing phenotypes, 2) karyotypic diversity among these phenotypic variants, 3) diversity in karyotypic stability, and 4) diversity in switching potential.

Adaptation through aneuploidization may be especially important for the survival of pathogenic microbes, which must endure fluctuating conditions within their host niches. This is supported by recent genomic surveys which have revealed pervasive aneuploidy in the genomes of pathogenic and clinical isolates of *S. cerevisiae*, *C. albicans*, and *C. neoformans*^40,45–47^. Moreover, aneuploidy has been associated with increased virulence^8,19^, thermotolerance^35^, and drug resistance^39^ in these fungal pathogens. Intriguingly, these pathogenic traits have also been associated with an isolate’s ability to develop different CMs, be it through a regulated CMR program, or through stochastic phenotype switching^8,18–20,48^. Although yet untested, it is tempting to speculate that aneuploid phenotypic variants such as those characterized in this study play key roles in pathogenic adaptation and persistence in a disease context. Future studies will focus on defining the contributions of aneuploidy-based phenotypic switches to population survival and adaptation, as well as how these switch events perpetuate genomic evolution and diversity over time.

---

**Figure S1.** Draft structure of Chr1^cir^. An annotated schematic generated using CLC-Genomics of the fusion site on Chr1^cir^ between a Ty element and the sub-telomeric sequences of the left telomere of Chr1 (top) and a draft assembly of the complete Chr1^cir^ molecule. Dark blue arrows denote genes; light blue arrows denote transposable elements; red and yellow regions denote telomeric sequences.

**Figure S2.** Structure of seg-CNAs identified in variant genomes. A-D. Cartoon schematics of translocations identified in the denoted variant genomes deduced from Illumina and Oxford Nanopore sequencing analysis combined with PFGE. Detailed descriptions of these Seg. CNAs are presented in Table S3.

**Figure S3.** Experimental design of the counter-selectable chromosome loss assay. A schematic illustrating the genotypic and phenotypic outcomes of selection for loss of Chr5^B^ in the YJM311 background. *CAN1* deletion, red box; Chr5^B^-located *CAN1* markers, yellow boxes.

## Experimental Methods

### Strain Construction

All experiments performed in this study were carried out using a clonal strain of the diploid clinical isolate YJM311, generously provided to us by Dr. Paul Magwene. To construct the strain used for the *CAN1* counterselection assays, a single copy of *CAN1* was deleted from Chr5A using a PCR product encoding the *HPHMX6* cassette, which confers resistance to the drug hygromycin B^49^. Correct targeting to and deletion of the *CAN1* locus was confirmed by PCR. Next, a PCR product consisting of *CAN1-KANMX* amplified from genomic DNA was integrated on the right arm of Chr5B at position 256375-257958 of the reference genome. Correct insertion or deletion of the *CAN1* was confirmed by PCR.

### Variant Recovery and Media

To isolate spontaneously arising complex or smooth phenotypic variants, YJM311 cells were streaked on solid YP^2.0%^ (10g/L yeast extract, 20 g/L Peptone, 20 g/L bacteriological agar, and 20g/L glucose (2.0%)) media and incubated at 30°C for 32 hours to allow single colonies to grow. Individual single colonies were each inoculated into 5mL liquid YP^2.0%^ cultures and incubated at 30°C for another 24 hours on a rotating drum. These cultures consisted of ~10^8^ cells, or approximately 30 generations derived from the original colony-forming cell. Cultures were diluted appropriately and plated such that ~200 individual cells were plated to either YP^2.0%^ or YP^0.5%^ (10g/L yeast extract, 20 g/L Peptone, 20 g/L bacteriological agar, and 5g/L glucose (0.5%)). Plates were incubated at 30°C for 3 days and then visually screened for colonies that displayed the variant phenotype. Total colonies were counted and the frequencies at which morphological variants appeared were calculated (Table S1). Candidate variant colonies were isolated, resuspended in water, spotted to fresh plates, and grown at 30°C for at least 3 days to confirm the persistence of the phenotypic variation. The entirety of each patch was then resuspended and frozen until DNA was prepared for genomic sequencing.

### Whole genome sequencing analysis

The genomes of 68 wild-type and variant clones were sequenced using Illumina short read whole genome sequencing and analyzed as described previously^23^. Briefly, Genomic DNA from each clone was isolated using the Yeastar Genomic DNA kit from Zymo Research. Pooled, barcoded libraries of individual genomes were generated using a Seqwell plexWell-96 kit. The final barcoded library was sequenced using an Illumina HiSeq sequencer. Illumina reads for each genome were mapped to the yeast reference genome (R64-2-1, yeastgenome.org) using CLC-Genomics software (Qiagen). Resulting read mapping files were then subjected to copy number and heterozygous single nucleotide polymorphism (hetSNP) variant analysis using the Nexus Copy Number software program (Biodiscovery). From this, we identified whole chromosome gains/losses, segmental duplications/deletions, and tracts of loss-of-heterozygosity (LOH). Aneuploidy was defined as the deviation in copy number of each individual homolog away from 1n. Using this definition, UPDs were scored as two CCNAs. SVs identified in Nexus were confirmed manually using CLC Genomics. The complete karyotypic analyses of variant clones are reported in Table S2. Several clones harboring chromosomal translocations (1^*SV*^, 22^*SV*^, 27^*SV*^) were further analyzed using Oxford Nanopore Technologies (ONT) Minion sequencing. High molecular weight DNA was prepared from cells, barcoded using the ONT ligation sequencing and native barcoding kits (SQK-LSK-109, EXP-NBD-104), and sequenced on a Minion flow cell (FLO-MIN106D). Reads were basecalled using ONT Guppy software and analyzed using CLC Genomics Workbench.

To investigate the mosaicism of Chr1 monosomy in our YJM311 stock and WT clones, we measured the log2 ratio of read coverage across a defined region of Chr1 (position 50,000-150,000). For this analysis, a log2 ratio of ~0 indicates that the chromosomal region is present at 2 copies, and a log2 ratio of −1 indicates that the chromosomal region is present at only one copy. Log2 ratio values that fall in between 0 and −1 indicate that the sample contains a mixture of cells harboring 1 or 2 copies of the defined genomic region. The log2 values, and corresponding copy number values for Chr1 are reported in Table S2. For comparison, we also measured the log2 ratio of read coverage at region on Chr7 (position 200,000-300,000) within these same genomes. Log2 values and corresponding copy number values for this region are also presented in Table S2 and demonstrate that copy number mosaicism in these clones is restricted to Chr1.

### Fluctuation Analysis

Cells from were streaked to YP2.0% and grown at 30°C for 48 hours to allow single colonies to form. Individual colonies were resuspended in 200uL 1x TE buffer (Thermofisher), diluted appropriately, and plated onto YP2.0% and canavanine-supplemented plates (20g/L glucose, 5g/L ammonium sulfate, 1.7g/L yeast nitrogen base without amino acids, 1.4g/L arginine dropout mix, 20g/L bacteriological agar, 0.6mg canavanine sulfate). Plates were incubated at 30°C for 72 hours after which colonies were counted. Colony count data were used to calculate rates and 95% confidence intervals using Flucalc, a MSS-MLE (Ma-Sandri-Sarkar Maximum Likelihood Estimator) calculator for Luria-Delbrück fluctuation analysis (flucalc.ase.tufts.edu)^50^. Calculated rates and confidence intervals of four independent plating experiments, in total equaling 24 independent assays, are presented in Table S3.

### Reversion sectoring assays

Cells from frozen stocks of clones 6^*SV*^, 99^*SV*^, 18^*SV*^, 28^*CV*^, 35^*CV*^, 31^*SV*^, and 17^SV^ were streaked to either YP^0.5%^ or YP^2.0%^ and grown at 30°C for 48 hours. Colonies completely displaying the variant phenotype were picked with a sterile toothpick and streaked to a fresh plate and grown for 48 hours, such that individual colonies could be resolved. Total colonies were counted and revertant sectors were quantified. Frequencies of reversions were calculated as the number of revertant sectors that arose in the total number of screened colonies and are presented in Table S4.

### Colony Imaging

Colonies were imaged using a Dino-lite Edge 1.3MP AF4115ZT polarized microscope and the companion DinoScope 2.0 software program. Images were processed using Adobe Photoshop, Premier Pro, and Premier Rush programs. For Fig. 1A, time-course imaging began when colonies had been grown for 48hrs.

### Pulsed Field Gel Electrophoresis

PFGE protocols and analysis were conducted as described previously^51,52^.

### Statistical Analysis

To determine if specific chromosomal aneuploidies were enriched in one class of variant clones relative to the other (complex variants vs. smooth variants), we used a Chi square goodness of fit test and Pearson residuals analysis to compare the distribution of observed frequencies of CCNAs for each chromosome. Pearson residual values outside the range of −1.96-1.96 (two standard deviations) indicated significant enrichment of the specific chromosome between complex and smooth clones. From this, we determined that CCNAs of Chr13 and Chr15 displayed a significant enrichment in complex and smooth clones respectively.

## Acknowledgements

We are grateful to Dr. Paul Magwene for sharing the strain YJM311. This study was supported by NIH/NIGMS awards 1K99GM13419301 to LRH and R35GM11978801 to JLA.

## Author Contributions

Conceptualization: LRH; Methodology: LRH; Investigation: LRH; Resources: LRH and JLA; Writing: LRH and JLA; Funding acquisition: JLA and LRH.

## Data and Materials Availability

Sequencing data for each clone in this study will be deposited on NCBI.

