## Supplemental Materials for "Systemic aneuploidization events drive phenotype switching in *Saccharomyces cerevisiae*"

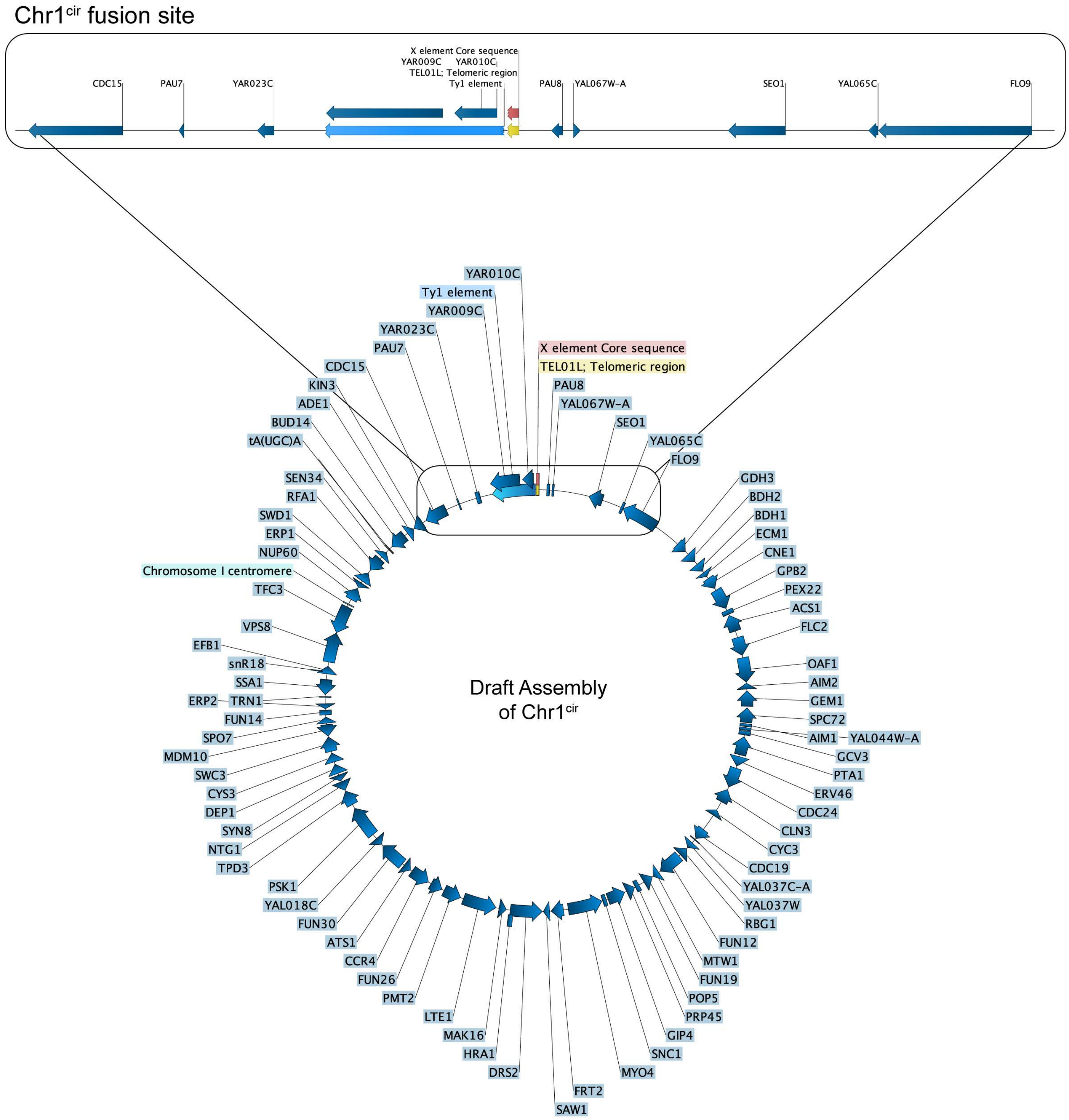


**Supplemental Figure 1**

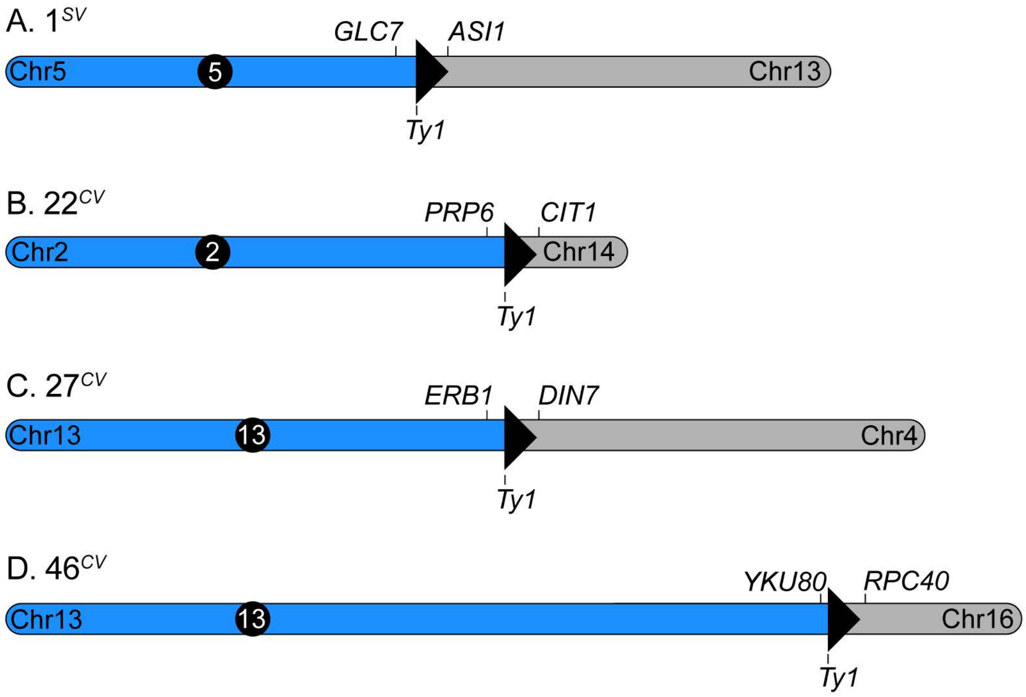


**Supplemental Figure 2**

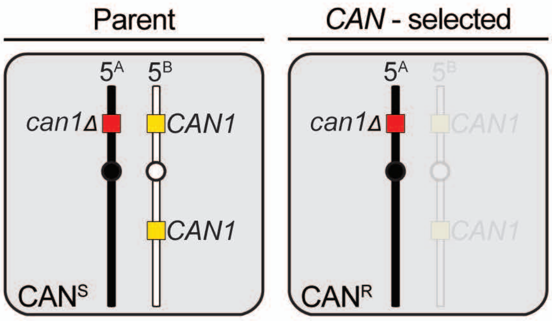


**Supplemental Figure 3**

| Table S1. Frequency calculations for the appearance of simple or complex variants | | |
|---|---|---|
| Frequency of complex colonies on YP-2.0% | | |
| Average frequency of variants | Number of Cultures | Total Number of Colonies |
| 1.59E-03 | 6 | 5648 |
| 1.66E-03 | 5 | 4896 |
| 9.92E-04 | 6 | 5454 |
| 9.26E-04 | 5 | 7344 |
| Frequency of simple colonies on YP-0.5% | | |
| Average frequency of variants | Number of Cultures | Total Number of Colonies |
| 1.59E-03 | 8 | 18893 |
| 4.06E-04 | 10 | 20185 |
| 9.63E-04 | 7 | 7421 |
| 9.82E-04 | 6 | 6088 |
| 1.18E-03 | 6 | 5501 |
| 9.32E-04 | 5 | 7325 |
| 1.29E-03 | 6 | 8742 |

Table S2. Copy Number Analysis of Chr1 in YJM311 and WT derivative clones. For comparison, copy number analysis for denoted region of Chr7 is also shown. Blue cells highlight WT clones that are monosomic for Chr1.

| | genomic region: chr1:50,000-150,000; 427 hetSNPs | | genomic region: chr7:200,000-300,000; 66 hetSNPs | |
|---|---|---|---|---|
| **Isolate** | **log2 coverage (median)** | **Copy Number** | **log2 coverage (median)** | **Copy Number** |
| YJM311 | -0.18 | 1.76 | -0.02 | 1.97 |
| 57 | -0.07 | 1.91 | -0.02 | 1.97 |
| 58 | -0.03 | 1.96 | -0.01 | 1.98 |
| 59 | -0.24 | 1.69 | -0.02 | 1.97 |
| 60 | -0.04 | 1.95 | -0.03 | 1.95 |
| 61 | -0.07 | 1.91 | -0.03 | 1.96 |
| 62 | -0.08 | 1.89 | -0.02 | 1.97 |
| 63 | -0.85 | 1.11 | -0.02 | 1.97 |
| 64 | -0.88 | 1.09 | -0.03 | 1.95 |
| 65 | -0.04 | 1.94 | -0.01 | 1.98 |
| 66 | -0.84 | 1.11 | -0.04 | 1.95 |
| 77 | -0.09 | 1.88 | -0.03 | 1.96 |
| 78 | -0.12 | 1.84 | -0.01 | 1.98 |
| 79 | -0.04 | 1.94 | -0.02 | 1.97 |
| 80 | -0.08 | 1.89 | -0.02 | 1.97 |
| 81 | -0.88 | 1.09 | -0.03 | 1.96 |
| 82 | -0.17 | 1.77 | -0.02 | 1.98 |
| 83 | -0.05 | 1.93 | -0.01 | 1.98 |
| 84 | -0.05 | 1.93 | -0.03 | 1.96 |
| 85 | -0.14 | 1.82 | -0.01 | 1.98 |
| 86 | -0.05 | 1.93 | -0.01 | 1.99 |

Table S3. Karyotypic analysis of variant genomes. A/B delineate each homolog in a pair. Bold cells indicate aneuploidies.

| Isolate Name | Variant Class | Chr1 | | **Chr2** | | Chr3 | | **Chr4** | | Chr5 | | **Chr6** | | Chr7 | | **Chr8** | | Chr9 | | **Chr10** | | Chr11 | | **Chr12** | | Chr13 | | **Chr14** | | Chr15 | | **Chr16** | | total # *de novo* SVs | Non-CCNA SVs |
|---|---|---|---|---|---|---|---|---|---|---|---|---|---|---|---|---|---|---|---|---|---|---|---|---|---|---|---|---|---|---|---|---|---|---|---|
| | | A | B | A | B | A | B | A | B | A | B | A | B | A | B | A | B | A | B | A | B | A | B | A | B | A | B | A | B | A | B | A | B | | |
| 1_SV | simple | 1 | 1 | 1 | 1 | 1 | 1 | 1 | 1 | 1 | 1 | 1 | 1 | 1 | 1 | 1 | 1 | 1 | 1 | 1 | 1 | 1 | 1 | 1 | 1 | 1 | 1 | 1 | 1 | 1 | 1 | 1 | 1 | 1 | Deletion of Chr5 (437267-Chr5RTel), Amplification of Chr13 (502144-Chr13RTel). |
| 2_SV | simple | **1** | **0** | 1 | 1 | 1 | 1 | 1 | 1 | 1 | 1 | 1 | 1 | 1 | 1 | 1 | 1 | 1 | 1 | 1 | 1 | 1 | 1 | 1 | 1 | 1 | 1 | 1 | 1 | **2** | **1** | 1 | 1 | 2 | |
| 3_SV | simple | **1** | **0** | 1 | 1 | 1 | 1 | 1 | 1 | 1 | 1 | 1 | 1 | 1 | 1 | **0** | **1** | 1 | 1 | 1 | 1 | 1 | 1 | 1 | 1 | 1 | 1 | 1 | 1 | 1 | 1 | 1 | 1 | 2 | |
| 4_SV | simple | **1** | **0** | 1 | 1 | 1 | 1 | 1 | 1 | 1 | 1 | 1 | 1 | 1 | 1 | 1 | 1 | 1 | 1 | **0** | **2** | 1 | 1 | 1 | 1 | 1 | 1 | 1 | 1 | 1 | 1 | 1 | 1 | 2 | |
| 5_SV | simple | 1 | 1 | 1 | 1 | 1 | 1 | **2** | **2** | 1 | 1 | 1 | 1 | 1 | 1 | 1 | 1 | 1 | 1 | **1** | **2** | 1 | 1 | **2** | **1** | **2** | **2** | 1 | 1 | 1 | 1 | 1 | 1 | 6 | |
| 6_SV | simple | **1** | **0** | 1 | 1 | 1 | 1 | 1 | 1 | **1** | **0** | 1 | 1 | 1 | 1 | 1 | 1 | 1 | 1 | 1 | 1 | 1 | 1 | 1 | 1 | 1 | 1 | 1 | 1 | 1 | 1 | 1 | 1 | 2 | |
| 15_SV | simple | 1 | 1 | 1 | 1 | 1 | 1 | 1 | 1 | 1 | 1 | 1 | 1 | 1 | 1 | 1 | 1 | 1 | 1 | **2** | **1** | 1 | 1 | 1 | 1 | 1 | 1 | 1 | 1 | **1** | **2** | 1 | 1 | 2 | |
| 17_SV | simple | 1 | 1 | 1 | 1 | 1 | 1 | **2** | **1** | 1 | 1 | 1 | 1 | **1** | **2** | 1 | 1 | 1 | 1 | 1 | 1 | 1 | 1 | 1 | 1 | 1 | 1 | 1 | 1 | 1 | 1 | 1 | 1 | 2 | |
| 18_SV | simple | 1 | 1 | 1 | 1 | **1** | **0** | 1 | 1 | 1 | 1 | 1 | 1 | 1 | 1 | 1 | 1 | 1 | 1 | 1 | 1 | 1 | 1 | 1 | 1 | 1 | 1 | 1 | 1 | 1 | 1 | 1 | 1 | 1 | |
| 19_SV | simple | 1 | 1 | 1 | 1 | 1 | 1 | 1 | 1 | 1 | 1 | 1 | 1 | 1 | 1 | 1 | 1 | 1 | 1 | 1 | 1 | 1 | 1 | 1 | 1 | 1 | 1 | 1 | 1 | **2** | **1** | 1 | 1 | 1 | |
| 20_SV | simple | 1 | 1 | **2** | **0** | **2** | **1** | 1 | 1 | **2** | **1** | **1** | **2** | **1** | **2** | 1 | 1 | **1** | **2** | **1** | **2** | **1** | **2** | **1** | **2** | **1** | **2** | **1** | **2** | **2** | **2** | **2** | **2** | 16 | |
| 29_SV | simple | 1 | 1 | 1 | 1 | 1 | 1 | **2** | **1** | 1 | 1 | 1 | 1 | **2** | **1** | 1 | 1 | 1 | 1 | **2** | **2** | 1 | 1 | 1 | 1 | 1 | 1 | 1 | 1 | **1** | **2** | 1 | 1 | 5 | |
| 30_SV | simple | 1 | 1 | 1 | 1 | 1 | 1 | **2** | **1** | 1 | 1 | 1 | 1 | 1 | 1 | 1 | 1 | **1** | **0** | 1 | 1 | 1 | 1 | 1 | 1 | 1 | 1 | 1 | 1 | 1 | 1 | 1 | 1 | 2 | |
| 31_SV | simple | 1 | 1 | 1 | 1 | 1 | 1 | 1 | 1 | 1 | 1 | 1 | 1 | 1 | 1 | 1 | 1 | 1 | 1 | 1 | 1 | 1 | 1 | 1 | 1 | 1 | 1 | 1 | 1 | **2** | **1** | 1 | 1 | 1 | |
| 32_SV | simple | 1 | 1 | 1 | 1 | 1 | 1 | 1 | 1 | 1 | 1 | **1** | **2** | 1 | 1 | 1 | 1 | 1 | 1 | 1 | 1 | 1 | 1 | 1 | 1 | **1** | **2** | 1 | 1 | **1** | **2** | 1 | 1 | 3 | |
| 33_SV | simple | 1 | 1 | 1 | 1 | 1 | 1 | 1 | 1 | **0** | **1** | 1 | 1 | 1 | 1 | 1 | 1 | 1 | 1 | 1 | 1 | 1 | 1 | 1 | 1 | 1 | 1 | 1 | 1 | 1 | 1 | 1 | 1 | 1 | |
| 34_SV | simple | 1 | 1 | **2** | **1** | 1 | 1 | **2** | **1** | 1 | 1 | 1 | 1 | 1 | 1 | 1 | 1 | **2** | **1** | **1** | **2** | 1 | 1 | 1 | 1 | 1 | 1 | **2** | **1** | 1 | 1 | **1** | **2** | 6 | |
| 43_SV | simple | 1 | 1 | 1 | 1 | 1 | 1 | 1 | 1 | 1 | 1 | 1 | 1 | 1 | 1 | 1 | 1 | 1 | 1 | 1 | 1 | 1 | 1 | 1 | 1 | 1 | 1 | 1 | 1 | **1** | **2** | 1 | 1 | 1 | |
| 44_SV | simple | 1 | 1 | **2** | **1** | 1 | 1 | 1 | 1 | 1 | 1 | 1 | 1 | 1 | 1 | 1 | 1 | **1** | **2** | 1 | 1 | 1 | 1 | 1 | 1 | 1 | 1 | 1 | 1 | **1** | **2** | 1 | 1 | 3 | |
| 50_SV | simple | 1 | 1 | 1 | 1 | 1 | 1 | 1 | 1 | 1 | 1 | 1 | 1 | 1 | 1 | 1 | 1 | 1 | 1 | 1 | 1 | 1 | 1 | 1 | 1 | 1 | 1 | 1 | 1 | **2** | **1** | 1 | 1 | 1 | |
| 51_SV | simple | 1 | 1 | 1 | 1 | 1 | 1 | 1 | 1 | 1 | 1 | 1 | 1 | 1 | 1 | 1 | 1 | 1 | 1 | 1 | 1 | 1 | 1 | 1 | 1 | 1 | 1 | 1 | 1 | **1** | **2** | 1 | 1 | 1 | |
| 52_SV | simple | 1 | 1 | **1** | **2** | 1 | 1 | 1 | 1 | 1 | 1 | **1** | **2** | **1** | **2** | 1 | 1 | **1** | **2** | 1 | 1 | 1 | 1 | **2** | **1** | 1 | 1 | 1 | 1 | **1** | **2** | 1 | 1 | 6 | |
| 97_SV | simple | **1** | **0** | 1 | 1 | 1 | 1 | 1 | 1 | **1** | **0** | 1 | 1 | 1 | 1 | 1 | 1 | 1 | 1 | 1 | 1 | 1 | 1 | 1 | 1 | 1 | 1 | 1 | 1 | 1 | 1 | 1 | 1 | 2 | |
| 98_SV | simple | 1 | 1 | 1 | 1 | 1 | 1 | 1 | 1 | 1 | 1 | 1 | 1 | 1 | 1 | **1** | **0** | 1 | 1 | 1 | 1 | 1 | 1 | 1 | 1 | 1 | 1 | 1 | 1 | 1 | 1 | 1 | 1 | 1 | |
| 99_SV | simple | 1 | 1 | 1 | 1 | 1 | 1 | 1 | 1 | 1 | 1 | 1 | 1 | 1 | 1 | 1 | 1 | **1** | **0** | 1 | 1 | 1 | 1 | 1 | 1 | 1 | 1 | 1 | 1 | 1 | 1 | 1 | 1 | 1 | |
| 102_SV | simple | 1 | 1 | 1 | 1 | 1 | 1 | **1** | **2** | 1 | 1 | 1 | 1 | **1** | **2** | 1 | 1 | 1 | 1 | 1 | 1 | 1 | 1 | 1 | 1 | 1 | 1 | 1 | 1 | 1 | 1 | 1 | 1 | 2 | |
| 22_CV | complex | 1 | 1 | 1 | 1 | 1 | 1 | 1 | 1 | 1 | 1 | 1 | 1 | 1 | 1 | 1 | 1 | 1 | 1 | 1 | 1 | 1 | 1 | 1 | 1 | 1 | 1 | 1 | 1 | 1 | 1 | 1 | 1 | 1 | Deletion of Chr2 (352193-Chr2RTel), Deletion of Chr14 (Chr14LTel-631061) |
| 23_CV | complex | 1 | 1 | 1 | 1 | 1 | 1 | 1 | 1 | 1 | 1 | 1 | 1 | 1 | 1 | 1 | 1 | 1 | 1 | **1** | **2** | 1 | 1 | 1 | 1 | **2** | **1** | 1 | 1 | 1 | 1 | 1 | 1 | 3 | |
| 24_CV | complex | 1 | 1 | 1 | 1 | 1 | 1 | **1** | **2** | 1 | 1 | 1 | 1 | 1 | 1 | 1 | 1 | 1 | 1 | 1 | 1 | 1 | 1 | 1 | 1 | **1** | **2** | 1 | 1 | 1 | 1 | 1 | 1 | 2 | |
| 25_CV | complex | 1 | 1 | 1 | 1 | 1 | 1 | 1 | 1 | 1 | 1 | 1 | 1 | 1 | 1 | 1 | 1 | **1** | **2** | 1 | 1 | 1 | 1 | 1 | 1 | **2** | **1** | 1 | 1 | **2** | **1** | 1 | 1 | 3 | |
| 26_CV | complex | 1 | 1 | 1 | 1 | 1 | 1 | 1 | 1 | 1 | 1 | 1 | 1 | 1 | 1 | 1 | 1 | 1 | 1 | 1 | 1 | 1 | 1 | 1 | 1 | **2** | **1** | 1 | 1 | 1 | 1 | 1 | 1 | 1 | |
| 27_CV | complex | 1 | 1 | 1 | 1 | 1 | 1 | 1 | 1 | 1 | 1 | 1 | 1 | 1 | 1 | 1 | 1 | 1 | 1 | 1 | 1 | 1 | 1 | 1 | 1 | 1 | 1 | 1 | 1 | 1 | 1 | 1 | 1 | 1 | Amplification of Chr4 (979209-Chr4RTel), Amplification of Chr13 (Chr13LTel-370517). |
| 28_CV | complex | **0** | **1** | 1 | 1 | 1 | 1 | 1 | 1 | 1 | 1 | 1 | 1 | 1 | 1 | 1 | 1 | 1 | 1 | **2** | **1** | 1 | 1 | 1 | 1 | 1 | 1 | 1 | 1 | 1 | 1 | 1 | 1 | 2 | |
| 35_CV | complex | 1 | 1 | 1 | 1 | 1 | 1 | 1 | 1 | 1 | 1 | 1 | 1 | 1 | 1 | 1 | 1 | 1 | 1 | 1 | 1 | 1 | 1 | 1 | 1 | **1** | **2** | 1 | 1 | 1 | 1 | 1 | 1 | 1 | |
| 36_CV | complex | **1** | **0** | 1 | 1 | 1 | 1 | 1 | 1 | 1 | 1 | **1** | **0** | 1 | 1 | 1 | 1 | 1 | 1 | **1** | **0** | 1 | 1 | 1 | 1 | 1 | 1 | 1 | 1 | 1 | 1 | **1** | **0** | 4 | |
| 37_CV | complex | **1** | **0** | 1 | 1 | 1 | 1 | 1 | 1 | 1 | 1 | 1 | 1 | 1 | 1 | 1 | 1 | 1 | 1 | 1 | 1 | 1 | 1 | 1 | 1 | **1** | **2** | 1 | 1 | 1 | 1 | 1 | 1 | 2 | |
| 38_CV | complex | 1 | 1 | 1 | 1 | 1 | 1 | 1 | 1 | 1 | 1 | 1 | 1 | 1 | 1 | 1 | 1 | 1 | 1 | 1 | 1 | 1 | 1 | 1 | 1 | **2** | **1** | 1 | 1 | 1 | 1 | 1 | 1 | 1 | |
| 39_CV | complex | 1 | 1 | 1 | 1 | 1 | 1 | 1 | 1 | 1 | 1 | 1 | 1 | 1 | 1 | 1 | 1 | 1 | 1 | 1 | 1 | 1 | 1 | 1 | 1 | **2** | **1** | 1 | 1 | 1 | 1 | 1 | 1 | 1 | |
| 40_CV | complex | **2** | **2** | 1 | 1 | 1 | 1 | 1 | 1 | 1 | 1 | **1** | **2** | **2** | **1** | **2** | **1** | **1** | **2** | **2** | **1** | 1 | 1 | 1 | 1 | 1 | 1 | 1 | 1 | 1 | 1 | **1** | **2** | 9 | |
| 41_CV | complex | **2** | **1** | 1 | 1 | 1 | 1 | 1 | 1 | 1 | 1 | 1 | 1 | 1 | 1 | 1 | 1 | 1 | 1 | 1 | 1 | 1 | 1 | 1 | 1 | **2** | **1** | 1 | 1 | 1 | 1 | **2** | **1** | 4 | |
| 42_CV | complex | 1 | 1 | 1 | 1 | 1 | 1 | 1 | 1 | 1 | 1 | 1 | 1 | 1 | 1 | 1 | 1 | 1 | 1 | 1 | 1 | 1 | 1 | 1 | 1 | **1** | **2** | 1 | 1 | 1 | 1 | **1** | **2** | 2 | |
| 46_CV | complex | 1 | 1 | 1 | 1 | 1 | 1 | 1 | 1 | 1 | 1 | 1 | 1 | 1 | 1 | 1 | 1 | 1 | 1 | 1 | 1 | 1 | 1 | 1 | 1 | 1 | 1 | 1 | 1 | 1 | 1 | 1 | 1 | 1 | Amplification of Chr13 (Chr13LTel-480490), Amplification of Chr16 (745828-Chr16RTel) |
| 47_CV | complex | 1 | 1 | 1 | 1 | 1 | 1 | 1 | 1 | 1 | 1 | 1 | 1 | **2** | **1** | 1 | 1 | 1 | 1 | **2** | **1** | 1 | 1 | 1 | 1 | 1 | 1 | 1 | 1 | 1 | 1 | 1 | 1 | 2 | |
| 48_CV | complex | 1 | 1 | 1 | 1 | 1 | 1 | 1 | 1 | 1 | 1 | 1 | 1 | 1 | 1 | 1 | 1 | 1 | 1 | 1 | 1 | 1 | 1 | 1 | 1 | **1** | **2** | 1 | 1 | 1 | 1 | 1 | 1 | 1 | |
| 49_CV | complex | 1 | 1 | 1 | 1 | 1 | 1 | 1 | 1 | 1 | 1 | 1 | 1 | 1 | 1 | **2** | **1** | 1 | 1 | 1 | 1 | 1 | 1 | 1 | 1 | 1 | 1 | 1 | 1 | 1 | 1 | 1 | 1 | 2 | Interstitial LOH at *ENA* array on Chr4 |
| 53_CV | complex | **1** | **0** | 1 | 1 | 1 | 1 | 1 | 1 | 1 | 1 | 1 | 1 | 1 | 1 | 1 | 1 | 1 | 1 | 1 | 1 | 1 | 1 | 1 | 1 | 1 | 1 | 1 | 1 | 1 | 1 | 1 | 1 | 1 | |
| 55_CV | complex | 1 | 1 | 1 | 1 | 1 | 1 | 1 | 1 | 1 | 1 | 1 | 1 | 1 | 1 | 1 | 1 | 1 | 1 | 1 | 1 | 1 | 1 | 1 | 1 | **1** | **2** | 1 | 1 | 1 | 1 | 1 | 1 | 1 | |
| 56_CV | complex | 1 | 1 | 1 | 1 | 1 | 1 | 1 | 1 | 1 | 1 | 1 | 1 | 1 | 1 | 1 | 1 | 1 | 1 | 1 | 1 | 1 | 1 | 1 | 1 | 1 | 1 | 1 | 1 | 1 | 1 | 1 | 1 | 0 | |
| 57_WT | wild type | 1 | 1 | 1 | 1 | 1 | 1 | 1 | 1 | 1 | 1 | 1 | 1 | 1 | 1 | 1 | 1 | 1 | 1 | 1 | 1 | 1 | 1 | 1 | 1 | 1 | 1 | 1 | 1 | 1 | 1 | 1 | 1 | 0 | |
| 58_WT | wild type | 1 | 1 | 1 | 1 | 1 | 1 | 1 | 1 | 1 | 1 | 1 | 1 | 1 | 1 | 1 | 1 | 1 | 1 | 1 | 1 | 1 | 1 | 1 | 1 | 1 | 1 | 1 | 1 | 1 | 1 | 1 | 1 | 0 | |
| 59_WT | wild type | 1 | 1 | 1 | 1 | 1 | 1 | 1 | 1 | 1 | 1 | 1 | 1 | 1 | 1 | 1 | 1 | 1 | 1 | 1 | 1 | 1 | 1 | 1 | 1 | 1 | 1 | 1 | 1 | 1 | 1 | 1 | 1 | 0 | |
| 60_WT | wild type | 1 | 1 | 1 | 1 | 1 | 1 | 1 | 1 | 1 | 1 | 1 | 1 | 1 | 1 | 1 | 1 | 1 | 1 | 1 | 1 | 1 | 1 | 1 | 1 | 1 | 1 | 1 | 1 | 1 | 1 | 1 | 1 | 0 | |
| 61_WT | wild type | 1 | 1 | 1 | 1 | 1 | 1 | 1 | 1 | 1 | 1 | 1 | 1 | 1 | 1 | 1 | 1 | 1 | 1 | 1 | 1 | 1 | 1 | 1 | 1 | 1 | 1 | 1 | 1 | 1 | 1 | 1 | 1 | 0 | |
| 62_WT | wild type | 1 | 1 | 1 | 1 | 1 | 1 | 1 | 1 | 1 | 1 | 1 | 1 | 1 | 1 | 1 | 1 | 1 | 1 | 1 | 1 | 1 | 1 | 1 | 1 | 1 | 1 | 1 | 1 | 1 | 1 | 1 | 1 | 0 | |
| 63_WT | wild type | **1** | **0** | 1 | 1 | 1 | 1 | 1 | 1 | 1 | 1 | 1 | 1 | 1 | 1 | 1 | 1 | 1 | 1 | 1 | 1 | 1 | 1 | 1 | 1 | 1 | 1 | 1 | 1 | 1 | 1 | 1 | 1 | 0 | |
| 64_WT | wild type | **1** | **0** | 1 | 1 | 1 | 1 | 1 | 1 | 1 | 1 | 1 | 1 | 1 | 1 | 1 | 1 | 1 | 1 | 1 | 1 | 1 | 1 | 1 | 1 | 1 | 1 | 1 | 1 | 1 | 1 | 1 | 1 | 0 | |
| 65_WT | wild type | 1 | 1 | 1 | 1 | 1 | 1 | 1 | 1 | 1 | 1 | 1 | 1 | 1 | 1 | 1 | 1 | 1 | 1 | 1 | 1 | 1 | 1 | 1 | 1 | 1 | 1 | 1 | 1 | 1 | 1 | 1 | 1 | 0 | |
| 66_WT | wild type | **1** | **0** | 1 | 1 | 1 | 1 | 1 | 1 | 1 | 1 | 1 | 1 | 1 | 1 | 1 | 1 | 1 | 1 | 1 | 1 | 1 | 1 | 1 | 1 | 1 | 1 | 1 | 1 | 1 | 1 | 1 | 1 | 0 | |
| 77_WT | wild type | 1 | 1 | 1 | 1 | 1 | 1 | 1 | 1 | 1 | 1 | 1 | 1 | 1 | 1 | 1 | 1 | 1 | 1 | 1 | 1 | 1 | 1 | 1 | 1 | 1 | 1 | 1 | 1 | 1 | 1 | 1 | 1 | 0 | |
| 78_WT | wild type | 1 | 1 | 1 | 1 | 1 | 1 | 1 | 1 | 1 | 1 | 1 | 1 | 1 | 1 | 1 | 1 | 1 | 1 | 1 | 1 | 1 | 1 | 1 | 1 | 1 | 1 | 1 | 1 | 1 | 1 | 1 | 1 | 0 | |
| 79_WT | wild type | 1 | 1 | 1 | 1 | 1 | 1 | 1 | 1 | 1 | 1 | 1 | 1 | 1 | 1 | 1 | 1 | 1 | 1 | 1 | 1 | 1 | 1 | 1 | 1 | 1 | 1 | 1 | 1 | 1 | 1 | 1 | 1 | 0 | |
| 80_WT | wild type | 1 | 1 | 1 | 1 | 1 | 1 | 1 | 1 | 1 | 1 | 1 | 1 | 1 | 1 | 1 | 1 | 1 | 1 | 1 | 1 | 1 | 1 | 1 | 1 | 1 | 1 | 1 | 1 | 1 | 1 | 1 | 1 | 0 | |
| 81_WT | wild type | **1** | **0** | 1 | 1 | 1 | 1 | 1 | 1 | 1 | 1 | 1 | 1 | 1 | 1 | 1 | 1 | 1 | 1 | 1 | 1 | 1 | 1 | 1 | 1 | 1 | 1 | 1 | 1 | 1 | 1 | 1 | 1 | 0 | |
| 82_WT | wild type | 1 | 1 | 1 | 1 | 1 | 1 | 1 | 1 | 1 | 1 | 1 | 1 | 1 | 1 | 1 | 1 | 1 | 1 | 1 | 1 | 1 | 1 | 1 | 1 | 1 | 1 | 1 | 1 | 1 | 1 | 1 | 1 | 0 | |
| 83_WT | wild type | 1 | 1 | 1 | 1 | 1 | 1 | 1 | 1 | 1 | 1 | 1 | 1 | 1 | 1 | 1 | 1 | 1 | 1 | 1 | 1 | 1 | 1 | 1 | 1 | 1 | 1 | 1 | 1 | 1 | 1 | 1 | 1 | 0 | |
| 84_WT | wild type | 1 | 1 | 1 | 1 | 1 | 1 | 1 | 1 | 1 | 1 | 1 | 1 | 1 | 1 | 1 | 1 | 1 | 1 | 1 | 1 | 1 | 1 | 1 | 1 | 1 | 1 | 1 | 1 | 1 | 1 | 1 | 1 | 0 | |
| 85_WT | wild type | 1 | 1 | 1 | 1 | 1 | 1 | 1 | 1 | 1 | 1 | 1 | 1 | 1 | 1 | 1 | 1 | 1 | 1 | 1 | 1 | 1 | 1 | 1 | 1 | 1 | 1 | 1 | 1 | 1 | 1 | 1 | 1 | 0 | |
| 86_WT | wild type | 1 | 1 | 1 | 1 | 1 | 1 | 1 | 1 | 1 | 1 | 1 | 1 | 1 | 1 | 1 | 1 | 1 | 1 | 1 | 1 | 1 | 1 | 1 | 1 | 1 | 1 | 1 | 1 | 1 | 1 | 1 | 1 | 0 | |

| Table S4. Fluctuation analysis derived rates of Chr5 loss in YJM311 background | | | | |
|---|---|---|---|---|
| Selection | Rate | Upper 95% difference | Lower 95% difference | Number of cultures |
| Chr5 | 6.43E-06 | 3.8977E-06 | 3.1631E-06 | 8 |
| Chr5 | 1.45E-05 | 9.6505E-06 | 7.6756E-06 | 6 |
| Chr5 | 8.85E-06 | 7.3552E-06 | 5.5397E-06 | 5 |
| Chr5 | 1.17E-05 | 8.8314E-06 | 6.8155E-06 | 5 |
| Overall Average | 1.03538E-05 | 7.4337E-06 | 5.79848E-06 | |

| Table S5. Reversion frequencies for specific YJM311 derived morphological variants. | | | | |
|---|---|---|---|---|
| Variant | Genomic Alteration | Total Colonies Counted | Revertant Sectors | Frequency |
| 6_SV | monosomy Chr5 | 1270 | 71 | 5.59E-02 |
| 18_SV | monosomy Chr3 | 1364 | 16 | 1.17E-02 |
| 99_SV | monosomy Chr9 | 1299 | 16 | 1.23E-02 |
| 28_CV | trisomy Chr10 | 2556 | 1 | 3.91E-04 |
| 35_CV | trisomy Chr13 | 2604 | 3 | 1.15E-03 |
| 31_SV | trisomy Chr15 | 1132 | 3 | 2.65E-03 |
| 17_SV | trisomy Chr4/Chr7 | 1634 | 6 | 3.67E-03 |
